# MiSiPi-Rna: an integrated tool for characterizing small regulatory RNA processing

**DOI:** 10.1101/2023.05.07.539760

**Authors:** Taiya Jarva, Jialin Zhang, Alex Flynt

## Abstract

RNA interference (RNAi) is mediated by small (20-30 nucleotide) RNAs that are produced by complex processing pathways. In animals, three main classes are recognized: microRNAs (miRNAs), small-interfering RNAs (siRNAs) and piwi-interacting RNAs (piRNAs). Understanding of small RNA pathways has benefited from genetic models where key enzymatic events were identified that lead to stereotypical positioning of small RNAs relative to precursor transcripts. Increasingly there is interest in using RNAi in non-model systems due to ease of generating synthetic small RNA precursors for research and biotechnology. Unfortunately, small RNAs are often rapidly evolving, requiring investigation of a species’ endogenous small RNAs prior to deploying an RNAi approach. This can be accomplished through small non-coding RNA sequencing followed by applying various computational tools; however, the complexity and separately maintained packages lead to significant challenges for annotating global small RNA populations. To address this need, we developed a simple and efficient R package (MiSiPi-Rna) which can be used to characterize pre-selected loci with plots and statistics, aiding researchers understanding RNAi biology specific to their target species. Additionally, MiSiPi-Rna pioneers several computational approaches to identifying Dicer processing to assist annotation of miRNA and siRNA.

## 1. Introduction

RNA interference (RNAi) is post-transcriptional gene regulatory mechanism found in most eukaryotes that regulates translation and/or suppresses viruses and mobile elements through sequence-dependent recruitment of Argonaute/Piwi proteins (Carthew, 2006). The three major RNAi modes in animals are microRNAs (miRNAs), short-interfering (siRNAs), and piwi-interacting RNAs (piRNAs). miRNAs and siRNAs are cleaved from double-stranded RNA (dsRNA) by an RNase III enzyme, such as Dicer, producing duplexed 21-24 nucleotide (nt) RNAs with 2nt 3’ overhangs (Elbashir et al., 2001). miRNAs are excised from short ∼70nt hairpin precursors, while siRNA are processed from longer dsRNAs (Zamore et al., 2000). The distinction continues as miRNAs and siRNAs load into different Ago proteins, at least in ecdysozoan invertebrates, where they have unrelated functions. Altogether different from miRNAs and siRNAs, piRNAs are derived from fragments of transcripts cleaved by Piwi proteins directed by a pre-existing pool of piRNAs (Lau et al., 2006; Ro et al., 2007). Cleaved RNAs can be processed by the Zucchini/Mito-PLD RNase leading to phased piRNAs from single-stranded transcripts, or in the “ping pong” cycle where reciprocal piwi proteins (i.e. Aubergine vs Argonaute 3 in *D. melanogaster*) coordinate piRNA production from shared substrate RNAs (le Thomas et al., 2014).

Exploitation of RNAi over the past two decades has enabled genetic manipulation for research and commercial products in pharmaceuticals and agroscience. Despite nearly twenty years of advancements in the applications of RNAi, navigating RNAi pathways for experimental design remains challenging (Joga et al., 2016; Willow et al., 2021). Another issue is that analysis of small RNA sequencing is based on libraries created from ligation and is distinct from other RNA sequencing approaches. While this provides an opportunity to explore RNA processing through characterization of endogenously occurring small RNA ends, it does require specialized computational approaches. Currently a variety of tools that use varying methods are available for small RNA assessment, but many are limited to specific classes of RNAs, rely on sequence similarity to previously annotated noncoding RNAs (ncRNA), or require non-trivial prior experience with computational environments (Barturen et al., 2014; Friedländer et al., 2012; Gebert et al., 2017; Geles et al., 2021; Han, Wang, Zamore, et al., 2015; Huang et al., 2007; D. Li et al., 2016; Rosenkranz & Zischler, 2012; K. Wang et al., 2014; X. Wang et al., 2005; Wu et al., 2011). Furthermore, there is significant gap in the availability of siRNAs-specific tools–the class of small RNA most often used in genetic technologies.

The tool we describe here: ‘MiSiPi-Rna’ (<u>mi</u>RNA, <u>si</u>RNA, <u>pi</u>RNA) is intended to enable small RNA characterization with only basic knowledge of computational tools and RNAi biology. The package is available to install via Github (https://github.com/stupornova33/MiSiPi-RNA). Nearly all required software components are contained in the package, aside from a precompiled RNAfold executable from the ViennaRNA package, which should be downloaded separately (Gruber et al., 2008). The minimum required input is a binary alignment mapping (BAM) file of small RNA sequencing reads and a 3-column BED file of regions of interest that might such as regions of high small RNA expression. For the siRNA and miRNA modules, the reference genome used for the alignment is also required as a function argument in order to incorporate structure predictions. The package utilizes several Bioconductor genomics packages, including RSamtools, Biostrings, and GenomicRanges for fast manipulation of sequence data (Lawrence et al., 2013; Morgan et al., 2022; Pages et al., 2022).

## 2. Results and Interpretations

Small RNA sequencing data should be prepared for the MiSiPi-Rna package with standard computational methods (for example, adaptor trimming and quality checking), followed by aligning small RNA reads to a reference genome using Bowtie or other alignment tools with appropriate arguments for small RNAs (Langmead & Salzberg, 2012). For example, supplying Bowtie with the options -a -m 200 -v0 will return alignments that contain no mismatches, while allowing some multi-mapping that can be controlled through the -m option. The method of identifying regions of interest will be specific to the needs of the user and the depth of the sequencing data, however, general processing with SAMTools and BEDTools can identify regions of high small RNA expression using mapped sequence reads and the reference genome (Supp Fig. 1A, Supplementary Methods) (H. Li et al., 2009; Quinlan & Hall, 2010).

The package contains modules for each class of small RNAs which can be run separately or all at once for holistic locus characterization. With some exceptions, read data is initially processed similarly in all modules. The package RSamtools is used in accessory functions (OpenBamFile, makeBamDF, getChrMinus, and getChrPlus) to read the BAM file and create separate data frames containing reads from either the plus or minus strand for the region of interest, removing reads outside of the expected size of small RNAs (18-32nt), and retaining the minimal amount of information that is needed for downstream processing (Morgan et al., 2022). Alignments are subsequently processed according to module-specific functions which leverage characteristics of biogenesis specific to each type of small RNA (Sup. Fig. 1B).

### 2.1 micro-RNAs/short hairpins

miRNAs are ∼22nt length sequences generated from short hairpin, ∼70nt structures (Lee, 2002). After excision of the hairpin by Drosha in the nucleus, the loop sequence is removed by Dicer, yielding a short segment of double-stranded RNA (Grishok et al., 2001; Hutvágner et al., 2001; Ketting et al., 2001). One strand of the RNA becomes the ‘guide’ strand and is biased to participate in inhibition of mRNA translation via the Argonaute protein containing RISC complex, while the ‘passenger’ strand is preferentially degraded (Khvorova et al., 2003; Schwarz et al., 2003). To identify this biogenesis mode, the miRNA module first provides a read size distribution plot, which in the case of miRNAs should show most reads aligning at the locus to be 20-24nt (Fig 1A). In addition, a density plot is provided that is colored coded based on size of reads to assess if miRNA sized reads are present (Fig 1B).

**Figure 1.**
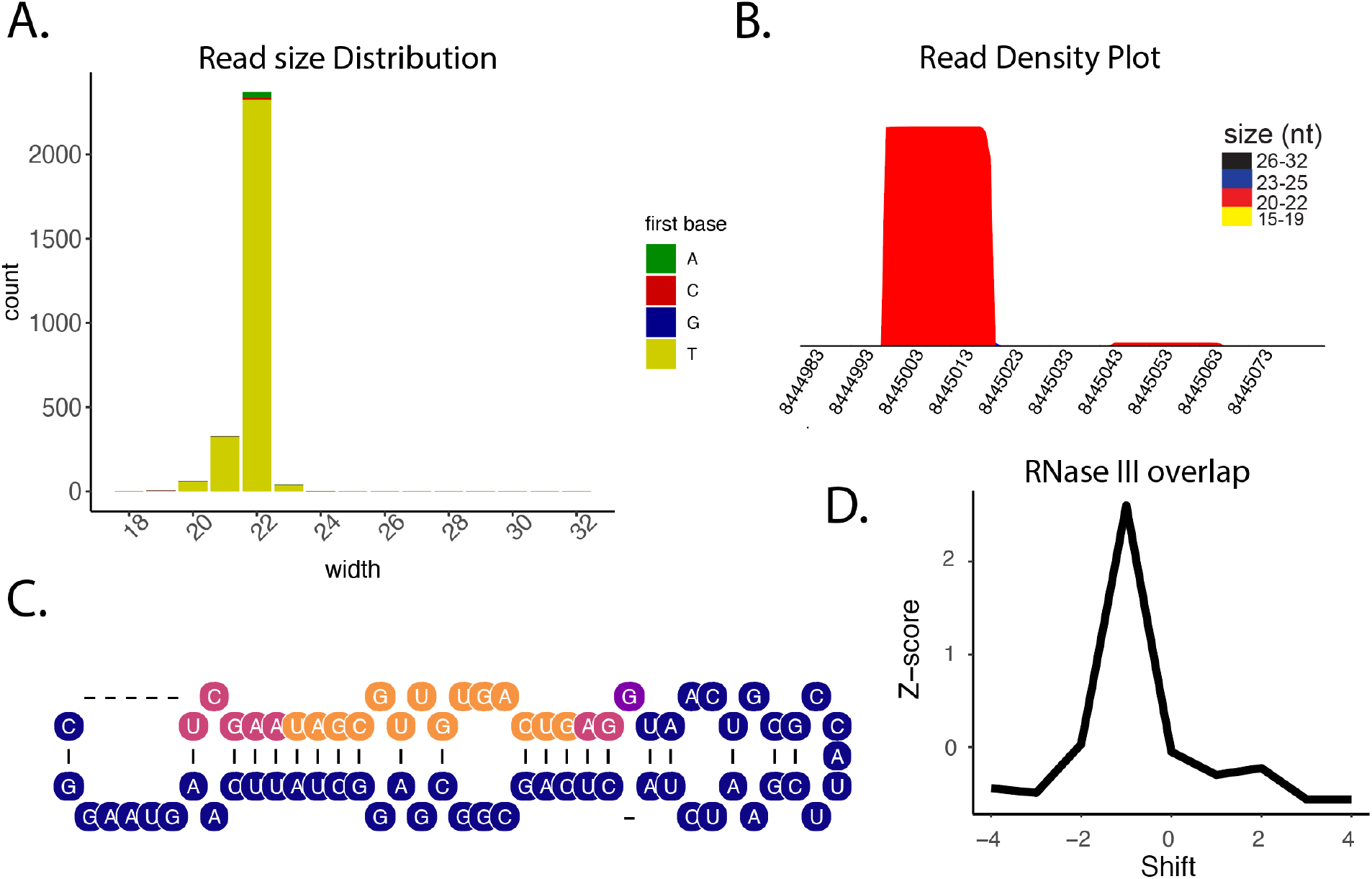
Example miRNA/shRNA output plots for *Drosophila melanogaster* mir-986. **A)** Read size distribution plot over the locus, colored by identity of first nucleotide. **B)** Read density over the locus. miRNAs typically have high read depth on the guide strand and low depth on the passenger strand. The color of the density plot is also colored by read size. **C**. Secondary structure plot colored by approximate expression of each nucleotide to assess overlay of expression with the precursor. **D**. Dicer probability score. Properly overhanging reads are counted over a range of shifted positions to calculate a probability score.

To further support positive identification of miRNAs, modules also take into consideration predicted structure. The package will output a secondary structure colored by relative expression which can be used to identify the location of major product (i.e. guide strand) that are derived from a hairpin (Fig. 1C). The final component of the miRNA module uses an algorithm to identify reads mapping within 60nt of each other on the same strand that potentially form miRNA guide-passenger duplexes. A major problem with annotation of miRNAs is the positive identification of RNase III processing, making automated characterization valuable to annotation efforts and may reduce pollution of incorrect annotations in public databases. During initial processing, a data frame is made of reads from a single strand, along with a copy of the data frame in which the 3’ ends of the reads are transformed by adding 60 nucleotides. The findOverlaps() function from the GenomicRanges package is then used to find artificially “overlapping” reads. The ends are then transformed back to their original positions and the genomic sequence between read pair ends is retrieved to represent a potential precursor hairpin (Sup. Fig. 2A). Overlapping hairpin arm designations are merged using the IRanges function reduce(), and followed by overlaying with RNAfold structure prediction from the ViennaRNA package. Probability of RNase III processing is calculated from the number of reads with the 2nt overhang in the context of the hairpin structure. This is followed by shifting the read start position from a range of -4 to 4 and again counting the number of reads with Dicer overhangs. If most reads are indeed RNase III-generated, the highest z-score will be found at a shift of 0 – frequent recovery of offset reads would show overhangs not characteristic of Drosha and Dicer processing (Fig. 1D, Sup. Fig. 2B).

### 2.2 Short-Interfering RNAs

siRNAs are short sequences (21-24nt) which like miRNAs are products of Dicer processing (Tijsterman et al., 2002). Two basic configurations are found for endogenous-siRNA (endo-siRNAs) precursors-double-stranded RNA (dsRNA) from intermolecular hybridization that could arise from bidirectional transcription, and long hairpin structures that fold into dsRNA. The siRNA module is designed to assist differentiation of siRNA types in addition to positive identification of Dicer processing found in all siRNA biogenesis. Initially, the module plots the read size distribution at the locus, which will show bias of 21-24nt reads for siRNAs, regardless of precursor type (Fig 2A). Next, since siRNAs are generated by Dicer in a processive manner, the module determines the probability of reads being juxtaposed tail to head or “phased” (Fig 2B) (Okamura, Chung, et al., 2008). For many Dicer substrates, initiation of cleavage appears to happen from heterogenous starting positions, which significantly complicates this analysis due to the overlaps of reads produced from different cleavage registers. To isolate a phasing signature the top 10% most abundant read alignments are used in the calculation. This represents the most common “paths” Dicer might take across the substrate. Lack of phasing signature for a candidate precursor may represent a challenge in isolating a Dicer path, or that there is an absence of a favored register of Dicer. Thus, no clear phasing does not explicitly rule out Dicer activity.

**Figure 2.**
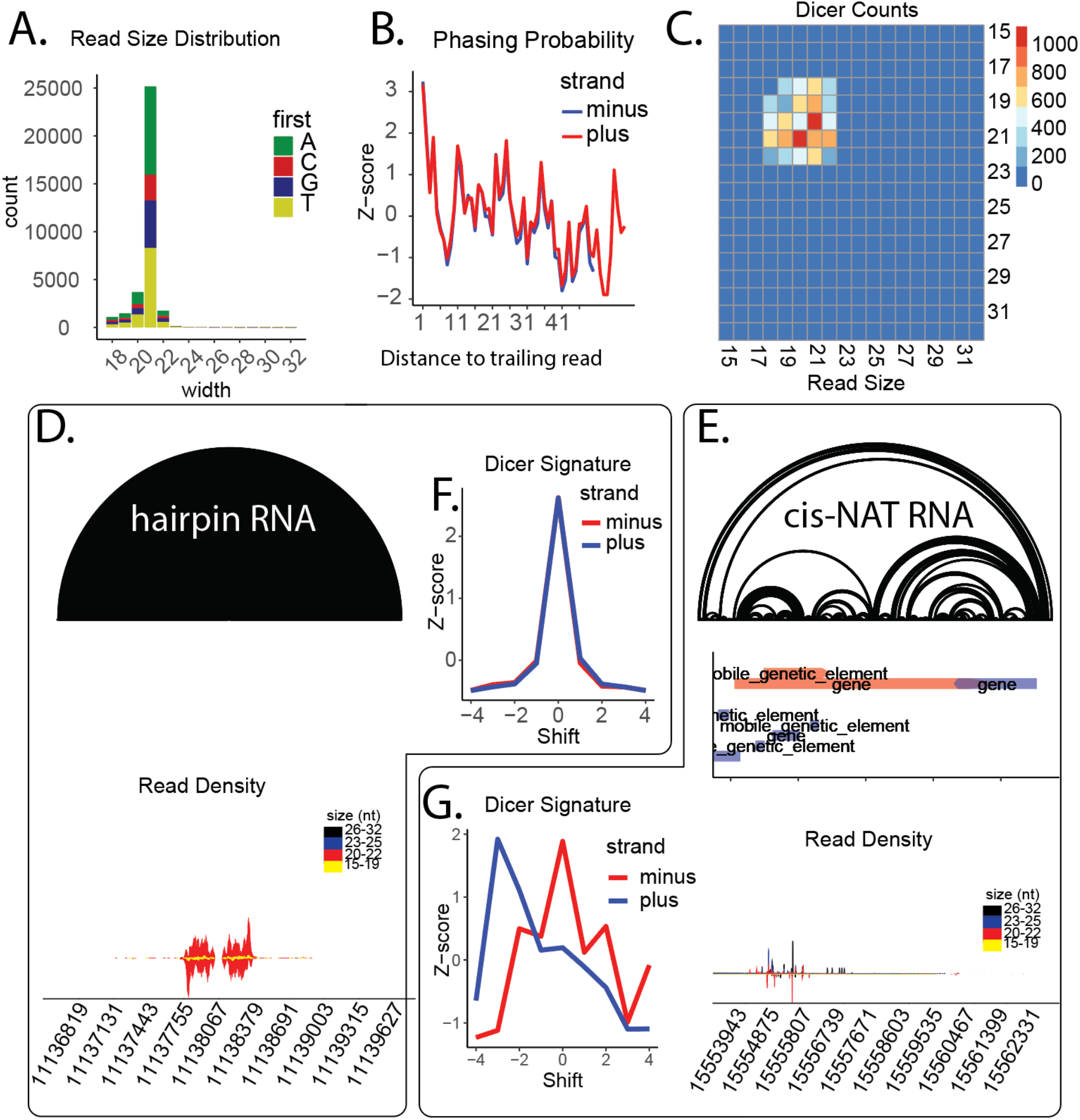
Endogenous siRNA characterization. **A**. Read size distribution over siRNA locus. **B**. Probability of reads mapping within n distance (phasing) of downstream reads as a z-score. **C**. Heatmap of reads which overlap and have a 3’ 2nt overhang which is characteristic of Dicer-2 processing. Heatmap was plotted using the ‘pheatmap’ package (Kolde, 2019) **D**. An arc plot generated for a hpRNA aligned with genomic region and density plot below colored by read size. **E**. An arc plot for a cisNAT region or hairpin region, genic features plotted below colored by strand (plus = red, minus = blue) and a density plot colored by size underneath. **F&G**. Dicer signature probabilities calculated across one strand based on structural predictions for a **(F)** hpRNA, as seen on left, and a **(G)** cis-NAT seen on right.

Next the package looks for reads that represent duplexed siRNAs with 2nt overhangs that are characteristic of Dicer processing. To achieve this, overlapping reads from both strands are examined for potential Dicer overlaps between all aligning pairs within the range. The product of this is a pairwise matrix of Dicer overlap counts that can visualized in a heatmap (Fig. 2C). siRNAs, regardless of precursor type, will likely show mapping of Dicer signature reads. This is the case even for many hairpin RNA-class siRNAs as arms of the structure are inverted repeats leading to reciprocal mapping artifacts to sense vs antisense strands.

The MiSiPi-Rna siRNA module provides additional, unique tools for examining siRNA precursor identity (Fig 2D-G). Fundamental to the approach is inclusion of secondary structure prediction by ViennaRNA, which is displayed as an arc diagram provided by the R4RNA and RRNA packages (Fig. 2D&E) (Bida & Maher, 2012; Lai et al., 2012). It is accompanied by an option for including genic or repetitive element annotations to identify feature overlaps. This is particularly useful for designating cis-natural antisense transcripts (cis-NATs) (Fig 2E) (Okamura, Balla, et al., 2008). The output also includes a density plot colored by read size that allows correlation of sites of siRNA production with structures. Finally, the siRNA module also provides a probability of Dicer processing from plus and minus genomic strands as well as across structural predictions. The module also calculates Dicer processing from a single strand by incorporating structural predictions to identify read pairs in a manner similar to the miRNA module (Fig 2F&G) Outputs of R4RNA include a table of paired base positions where a nucleotide ‘i’ is paired with a nucleotide ‘j’. The intervals are collapsed into sequential ‘i’ ranges and ‘j’ ranges greater than 15nt in length and allowing for up to 4 unpaired nucleotides. FindOverlaps() is used to create a set of reads which overlap with the ‘i’ ranges and a set of reads which overlap with the ‘j’ ranges. The read starts are then translated to their paired positions and the new read end is calculated from the start position and the width. From the overlapping uni-strand pairs, the dicer signature is calculated as well as shifted -4 to 4 as controls. For hairpin RNA precursors a strong Dicer signature can be observed (Fig 2F), while this is not the case for a bidirectionally transcribed cis-NAT (Fig 2G).

### 2.3 Piwi-Interacting RNAs

Biogenesis of piRNAs is wholly distinct from miRNA and siRNA as it is Dicer independent. This is immediately evident from the larger, ∼27-30nt size of piRNAs, which can be seen in the size distribution and density plots (Fig. 3A-B). Another feature of piRNAs is a bias for 1U residues (Beyret et al., 2012). Thus, when MiSiPi-Rna calculates size distribution for piRNAs it determines the proportion that are 1U (Fig. 3A). The package also supports functions for characterization of the two distinct piRNA biogenesis pathways: ping-pong and phasing. In both pathways, piRNAs are produced from transcripts targeted by pre-existing piRNAs, which permits identification of cleavage mode form sequencing data.

**Figure 3.**
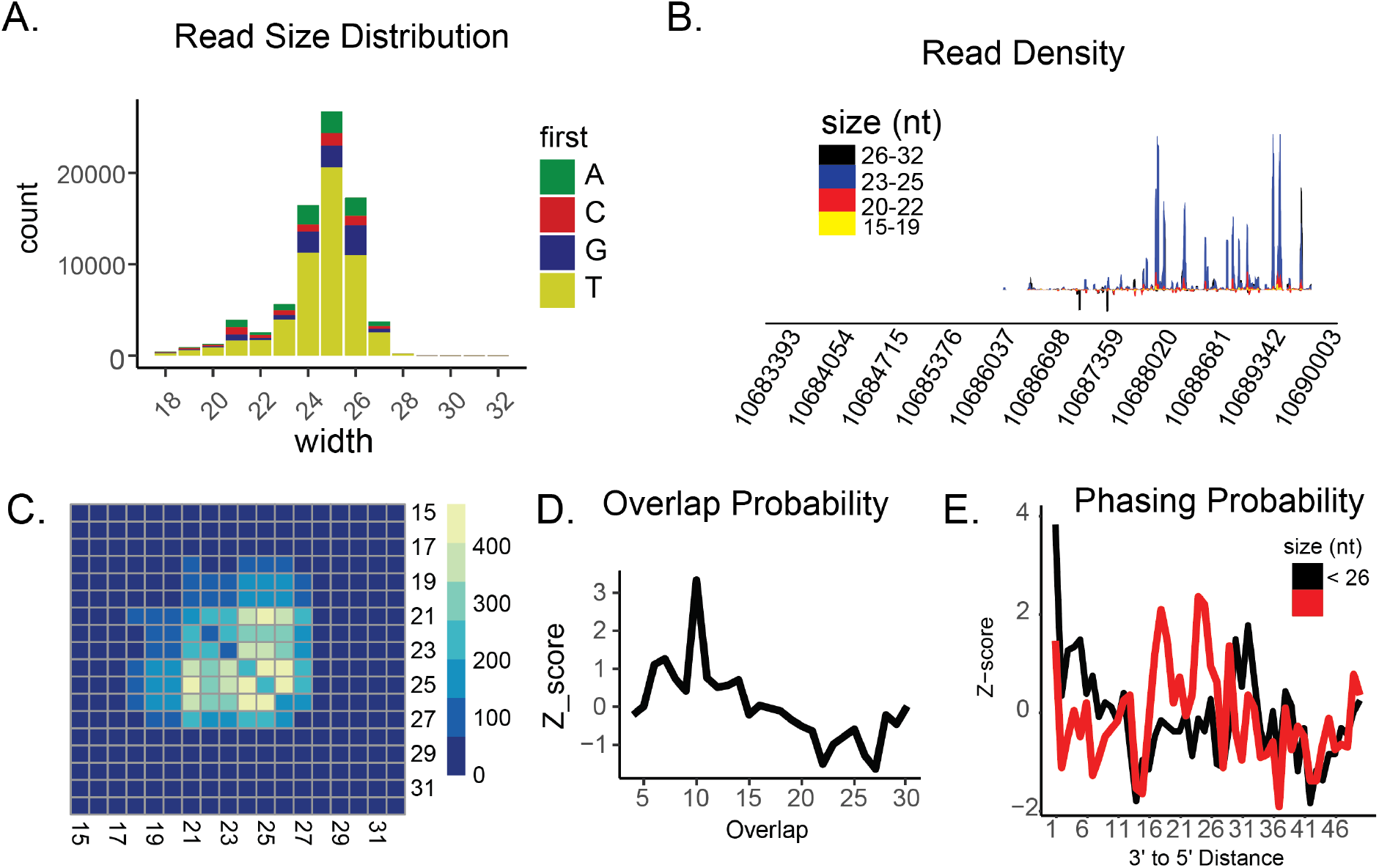
Example piRNA plots. **A**. Read size distribution over piRNA locus. **B**. Heatmap of reads which overlap by 10 nt plotted by read size. **C**. Probability plot for each overlap size calculated as a z-score. **D**. Read density over locus by read size. **E**. Probability of reads mapping within n nucleotides of downstream reads (phasing). The black line represents reads shorter than 26 nucleotides while red represents reads longer than 26 nucleotides.

Generation of piRNAs in response to cleavage guided by existing piRNAs is perhaps best captured in the ping-pong cycle. Here partner piwi proteins (i.e. Aubergine and Argonaute3 in *Drosophila*) cleave complementary strands, which then load and after resection become new piRNAs. Small RNAs generated via the ping-pong cycle overlap on their 5’ end of by 10 nt due to this reciprocal activity of piwi proteins (Sup. Fig. 2B). Thus, the module for piRNAs uses the GenomicRanges function findOverlaps to identify plus and minus strand reads that overlap by 10 nt for all alignments to the region. This is displayed as a count matrix of overlapping pairs (Fig. 3B). Shifts from the center show asymmetry in piRNAs partners and may reflect differences in the binding pocket size of their respective piRNA proteins (Han, Wang, Li, et al., 2015). The module also calculates the overlap length probability between read pairs, which is plotted as a simple line plot (Fig. 3C)

The other biogenesis mode is a type of phasing, which is initiated when a piRNA cleaves a transcript that becomes a substrate for the ribonuclease Zucchini/mito-PLD ((Brennecke et al., 2007; Mohn et al., 2015). It can be observed as conversion of single-stranded RNAs, that in some examples can be quite long (Nishida et al., 2007; Vagin et al., 2006). Phasing, as used to examine Dicer processing of siRNAs, is measured by calcuating the probability of reads being generated tail to head (Fig. 3C) (Gunawardane et al., 2007). The phased piRNA module uses a transformation method similar to the siRNA module, but only uses reads beginning with 1U (Sup. Fig. 2C). A phasing signature can be recognized when the distance is 1-3 nt apart, due to trimming of reads after initial cleavage by Zucchini/mito-PLD. The module will output a phasing probability plot, where the z-score is plotted separately for reads shorter than 26nt and reads 26nt or longer (Fig. 3E).

## Conclusions

While many sRNA annotation tools are available, there is a benefit to integrated software which can characterize all sRNA classes. This may increase deployment of RNAi for in recently sequenced or unannotated species. Indeed, while small RNA biology is well characterized in model species studies highlight the diversity of biogenesis between even related organisms (Flynt, 2021; Mondal et al., 2021) (Khanal et al., 2022). Notably, MiSiPi-RNA is one of the only tools available thus far which can classify siRNA loci. To facilitate overall small assessment the final function of MiSiPi-Rna is to output tables of values for phasing (both for siRNA and piRNA), dicer signatures (dual strand and across a hairpin), and pingpong signatures. Such tables are amendable to plotting with heatmaps and for clustering loci (Khanal et al., 2022).

Finally, MiSiPi-Rna is designed to be computationally efficient even over thousands of loci and on high-depth sequencing datasets and can be run on a laptop equipped with 8GB of RAM (Sup. Fig. 3). The package achieves this by being conscious of R language’s memory usage idiosyncrasies. For example, the package reads only one chromosome of data from the BAM file at once, extracting only reads from the region of interest, and retaining the minimal amount of data needed, e.g. the first letter of the nucleotide sequence and the start/stop positions instead of the entire sequence, which reduces the amount of data that is kept in memory. In addition, it uses fast sequence manipulation packages such as GenomicRanges and Biostrings, and the most computationally complex functions are written in C++ through the Rcpp package (Eddelbuettel & François, 2011). These approaches together also yield rapid computation such that 1000’s of loci can be assessed in compute processes that don’t take multiple days to complete (Sup Fig 3). The package modules have been validated here using annotated loci in *Drosophila* with publicly available datasets. Fundamentally however, MiSiPi-Rna can be immediately deployed in species for which a reference genome and small RNA sequencing data is available.

## Acknowledgments

AF and TJ are supported by NSF 1845978, and Mississippi INBRE: funded by an Institutional Development Award (IDeA) from the National Institute of General Medical Sciences of the National Institutes of Health under grant number P20GM103476. Alignment and processing of small RNA datasets was performed on Magnolia HPC at the University of Southern Mississippi, which is supported by the National Science Foundation under the Major Research Instrumentation (MRI) program via Grant # ACI 1626217.

