## supplemental_figures_methods for "MiSiPi-Rna: an integrated tool for characterizing small regulatory RNA processing"

### Supplement

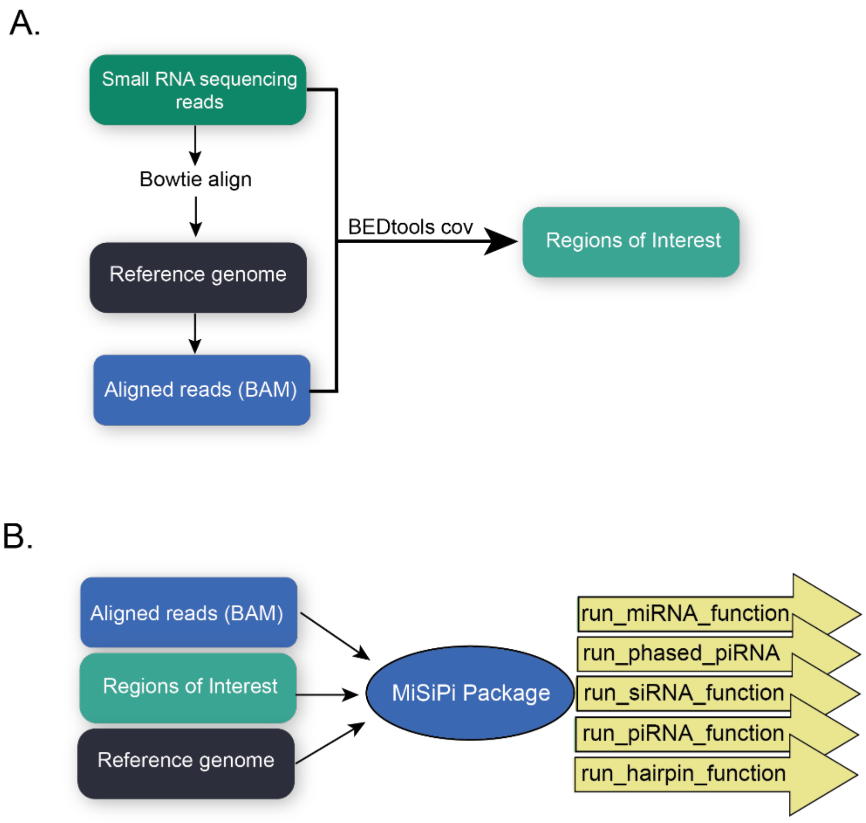


**Supplemental Figure 1**. Example MiSiPi Workflow. **A.** Raw sequence reads are aligned to a reference genome with Bowtie, giving a Binary Alignment Mapping file of aligned reads. The raw reads and the BAM file are used with BEDTools programs to identify regions of high small RNA expression. **B.** The BAM file, reference genome, and file containing regions of interest are provided as input to MiSiPi. The basic function commands for each module are shown.

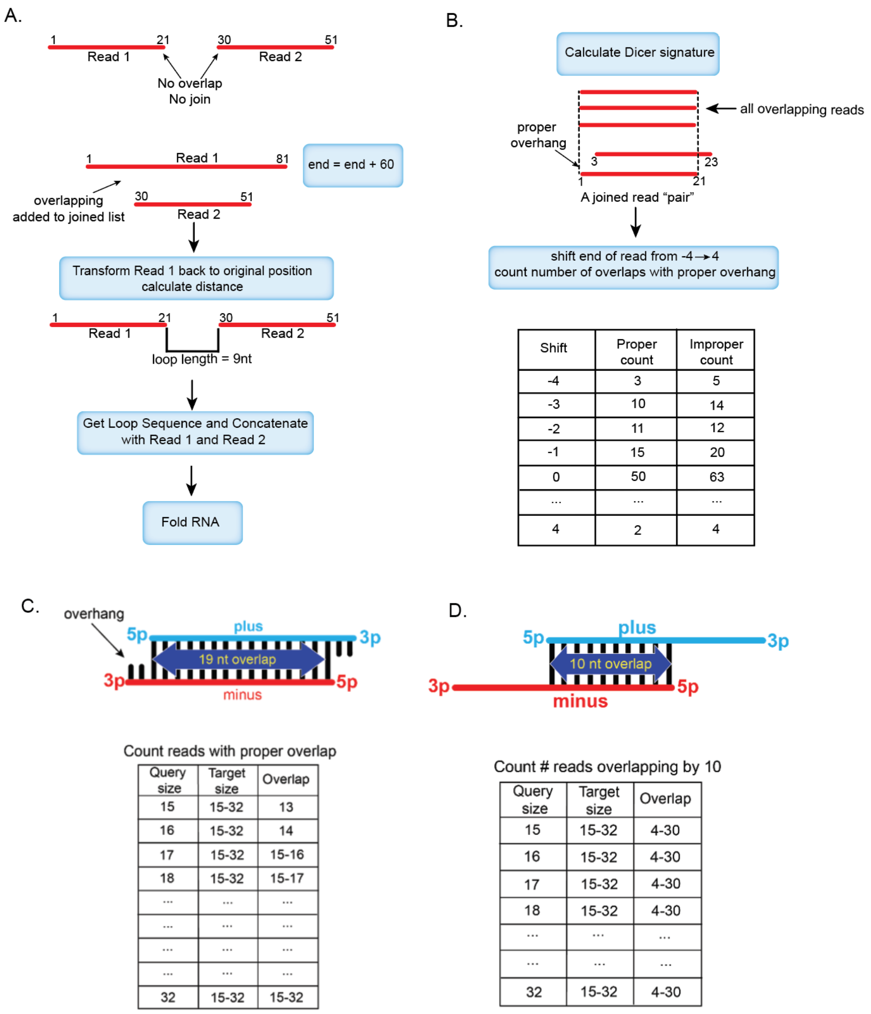


**Supplementary Figure 2**. **MiSiPi-RNA algorithm overviews. A.** The first step of the miRNA module is to identify reads mapping within 60nt of each other on the same strand. This is achieved by duplicating the data frame of reads from a single strand and incrementing the end of one set of reads by 60. Following this, the GenomicRanges function findOverlaps is used to identify 'overlapping' pairs. The read end positions are transformed to the original position and the sequence between read pairs is extracted from the genome. The concatenated sequence is then folded by RNAfold. The first steps of this algorithm are also used to test phasing in uni-strand piRNAs. **B.** The dicer probability is calculated by counting the number of reads overlapping each individual read that have a proper 2-nt overhang. The position of the read is shifted downstream or upstream by a range of -4 to 4, and the number of overlapping reads with a 2-nt overhang are re-counted. The z-score is calculated from the ratios of "proper" to "improper" overhangs. **C.** siRNA pairs are identified as reads which overlap between separate strands by approximately 19 nucleotides and which contain a 3' 2 nucleotide overhang. To generate the heatmap of overlaps (Fig. 2B), query reads ranging from 15-32nt are compared with overlapping reads ranging from 15-32nt. The pairs are tested for "proper" overlaps, (2 nucleotides less than the length of the query read) and the number of proper overlaps are counted. **D.** Similar to the siRNA module, piRNA pairs are identified as reads which overlap between both strands by 10 nucleotides, and a range of query, target sizes, and overlaps are used to plot the read sizes which overlap by 10nt (Fig. 3B).

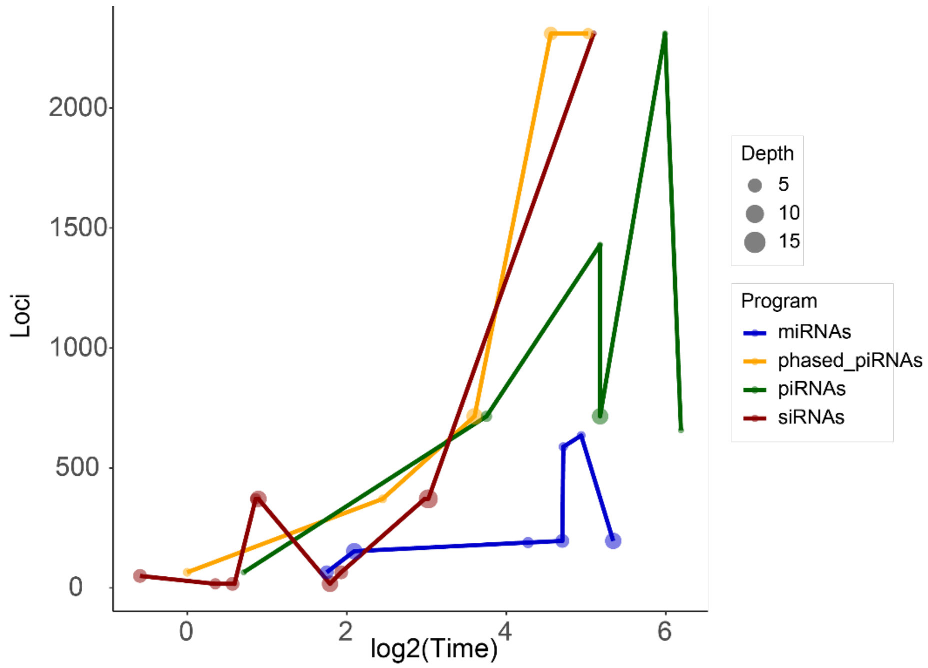


**Supplementary Figure 3. Program run times by number of loci and sequencing depth.** Each module of the MiSiPi package was run on publicly available datasets and plotted by sequencing depth. Time recorded is in log(minutes).

*Supplementary Methods*

### Identifying small RNA regions of interest

#### miRNAs & siRNAs: get reads of only miRNA and siRNA length

```
awk 'BEGIN {OFS = "\n"} {header = $0; getline seq; getline qheader ; getline
qseq ; if (length(seq) >= 19 && length(seq) <= 23) {print header, seq,
qheader, qseq}}' \
< trimmed.fq > small.fastq
```

#### piRNAs: get reads of piRNA length

```
awk 'BEGIN {OFS = "\n"} {header = $0; getline seq; getline qheader ; getline
qseq ; if(length(seq) >= 23 && length(seq) <= 30) {print header, seq,
qheader, qseq}}' \
< trimmed.fq > large.fastq
```

#### Realign reads to genome

```
bowtie -p 10 -a -m100 --best --strata --no-unal genome.fna small.fastq -S |
samtools view -@ 10 -q 10 -b |samtools sort -@ 10 -m 6G > small.bam
samtools index small.bam

bowtie -p 20 -a -m100 --no-unal genome.fna large.fastq -S | samtools view -@
10 -q 10 -b | samtools sort -@ 10 -m 6G > large.bam
samtools index large.bam
```

Calculate coverage of all reads over genome

```
bedtools genomecov -bg -ibam all.bam | awk '$4 > 100' > HE.tmp.bedgraph
```

Get large regions of high expression

```
bedtools merge -d 500 -i HE.tmp.bedgraph > HE.tmp.merge.bed

awk '{n=$2; x=$3; print $1"\t"$2"\t"$3"\t"x-n}' < HE.tmp.merge.bed |
awk '$4 > 40' > HE.all.bed #all highly expressed
```

Get potential regions of high miRNA/siRNA RNA expresssion

```
bedtools multicov -bams small.bam all.bam -bed HE.all.bed |
awk '{n=$5; x=$6; print $1"\t"$2"\t"$3"\t"n/x}' |
awk '$4 > 0.5' > HE.small.bed # only high expressed 21-23 nt
```

Get potential regions of high piRNA expression

```
bedtools multicov -bams large.bam all.bam -bed HE.all.bed |
awk '{n=$5; x=$6; print $1"\t"$2"\t"$3"\t"n/x}' | awk '$4 > 0.5' >
HE.large.bed #only high expressed 23-30 nt
```
